# The *Ralstonia pseudosolanacearum* effector RipE1 is recognized at the plasma membrane by *NbPtr1, Nicotiana benthamiana* homolog of *Pseudomonas tomato race 1*

**DOI:** 10.1101/2023.03.14.532688

**Authors:** Boyoung Kim, Injae Kim, Wenjia Yu, Haseong Kim, Ye Jin Ahn, Kee Hoon Sohn, Alberto P. Macho, Cécile Segonzac

**Affiliations:** Department of Agriculture, Forestry and Bioresources, Seoul National University, Seoul 08826, Republic of Korea; Plant Immunity Research Center, Seoul National University, Seoul 08826, Republic of Korea; Shanghai Center for Plant Stress Biology, CAS Center for Excellence in Molecular Plant Sciences, Chinese Academy of Sciences, Shanghai 201602, China; Department of Life Sciences, Pohang University of Science and Technology, Pohang 37673, Republic of Korea; Department of Agricultural Biotechnology, Seoul National University, Seoul 08826, Republic of Korea; Plant Genomics and Breeding Institute, Seoul National University, Seoul 08826, Republic of Korea; Research Institute of Agriculture and Life Sciences, Seoul National University, Seoul 08826, Republic of Korea

**Keywords:** Type III secreted effectors, plant immune system, nucleotide-binding leucine-rich repeat receptor, virus-induced gene silencing, subcellular localization

## Abstract

The bacterial wilt disease caused by soil-borne bacteria of the *Ralstonia solanacearum* species complex (RSSC) threatens important crops worldwide. Only a few immune receptors conferring resistance to this devastating disease are known so far. Individual RSSC strains deliver around 70 different type III secretion system effectors into host cells to manipulate the plant physiology and dampen immune responses. RipE1 is an effector conserved across RSSC isolated from diverse plant species and triggers immune responses in the model Solanaceae *Nicotiana benthamiana*. Here, we used multiplexed virus-induced gene silencing of the nucleotide-binding and leucine-rich repeat receptor family to identify the genetic basis of RipE1 recognition in *N. benthamiana*. Specific silencing of the *N. benthamiana* homolog of *Solanum lycopersicoides Pseudomonas tomato race 1* gene (*NbPtr1*) completely abolished RipE1-induced hypersensitive response and immunity to *Ralstonia pseudosolanacearum*. In Nb*-ptr1* knock-out plants, expression of the native *NbPtr1* coding sequence was sufficient to restore RipE1 recognition. In addition to the putative catalytic triad Cys-His-Asp, RipE1 association with the host cell plasma membrane was found necessary for NbPtr1-dependent recognition. Furthermore, we found that NbPtr1-dependent recognition of RipE1 natural variants is polymorphic suggesting the coevolutionary nature of this interaction. This work hence provides an additional evidence for the indirect mode of activation of NbPtr1 and supports NbPtr1 relevance for resistance to bacterial wilt disease in Solanaceae.

## Main text

Bacterial wilt is a devastating disease threatening the production of diverse crops worldwide. The causal agent belongs to the *Ralstonia solanacearum* species complex (RSSC) which consists of phylotypes from different geographic origins. RSSC was recently recategorized into three different species *R. solanacearum, R. pseudosolanacearum* and *R. syzygii* (Fegan and Prior, 2005; Prior et al., 2016; Wicker et al., 2012). These soil-borne bacteria enter the plant body through wound or natural openings to reach the vasculature where exponential multiplication and exopolysaccharide production eventually block the water flow leading to typical wilting symptoms (Lowe-Power et al., 2018; Mansfield et al., 2012). Compared to other bacterial phytopathogens, individual RSSC isolates deploy an unusually large number of type III secretion system (T3S) effectors that collectively contribute to pathogenicity and virulence (Landry et al., 2020; Peeters et al., 2013). After delivery into the host cell, T3S effectors function to suppress defense and/or manipulate the host metabolism (Macho 2016; Toruño et al., 2016).

Although comparisons of effector repertoire across RSSC isolates revealed a large variation, around 30 effector families are broadly present and considered as “core effectors” (Peeters et al., 2013; Sabbagh et al., 2019). One such core effector, RipE1, is conserved in *R. pseudosolanacearum, R. solanacearum* and *R. syzygii* strains isolated from diverse plant species (Mukaihara et al., 2010; Sabbagh et al., 2019). RipE1 sequence bears homology with *Pseudomonas syringae* and *Xanthomonas* spp. effectors belonging to AvrPphB/HopX family and harbors the conserved cysteine-histidine-aspartic acid catalytic triad and domain A (Gimenez-Ibanez et al., 2014; Nimchuk et al., 2007; Sang et al., 2020). RipE1 transient expression in *Nicotiana benthamiana* suppresses plant defenses, such as the production of reactive oxygen species and this requires the presence of putative catalytic cysteine (C172) and domain A (Nakano et al., 2019; Sang et al., 2020). However, similar to other members of the AvrPphB/HopX family, RipE1 expression also induces a robust cell death, indicative of its recognition by the plant immune system (Jeon et al., 2020; Mansfield et al., 1994; Nimchuk et al., 2007; Sang et al., 2020). Consistently, RipE1 expression leads to the induction of defense-associated gene expression, accumulation of salicylic acid and reduced susceptibility to *R. pseudosolanacearum* in *N. benthamiana* and *Arabidopsis thaliana* (Sang et al., 2020).

The plant immune system relies on arrays of cell surface-localized and intracellular receptors that function in a network sensing non-self or modified-self molecules that result in defense activation and pathogen growth restriction (Jones and Dangl, 2006; Ngou et al., 2022). Intracellular immune receptors from the nucleotide-binding and leucine-rich repeat receptor (NLR) family are activated by binding to corresponding effectors or, more commonly, by monitoring the effector-directed modifications of host proteins (Kourelis and van der Hoorn, 2018). Remarkably, only a handful of immune receptors have been demonstrated to contribute to RSSC effector recognition. The atypical Toll-interleukin Resistance-NLR (TNL) RESISTANCE TO RALSTONIA SOLANACEARUM 1 (RRS1) recognizes the effector RipP2 in *Arabidopsis thaliana* (Deslandes et al., 1998, 2002). RRS1 harbors a WRKY C-terminal extension, which is acetylated by RipP2 (Le Roux et al., 2015; Sarris et al., 2015). In *Nicotiana* spp., the TNL RESISTANCE TO XOPQ1 (Roq1) recognizes *X. euvesicatoria* effector XopQ and its homologs in *P. syringae* (HopQ1) and *R. pseudosolanacearum* (RipB) through direct binding on Roq1 LRR domain (Martin et al., 2020; Nakano and Mukaihara, 2019; Schultink et al., 2017). Last to date, the *Solanum lycopersicoides* Coiled-Coil NLR (CNL) PSEUDOMONAS TOMATO RACE 1 (Ptr1) recognizes the *P. syringae* effector AvrRpt2 and its homolog in *R. pseudosolanacearum* RipBN (Mazo-Molina et al., 2019, 2020).

Ptr1 activation mechanism is not yet fully understood but evidence suggests that Ptr1 monitors the state of the plasma membrane-associated RPM1-INTERACTING PROTEIN 4 (RIN4), which is cleaved in presence of the cysteine-protease effectors AvrRpt2 or RipBN (Axtell and Staskawicz, 2003; Mackey et al., 2003; Mazo-Molina et al., 2020; Takemoto and Jones, 2005). In line with this hypothesis, the *N. benthamiana* homolog of *Ptr1* (*NbPtr1*) was recently shown to contribute to the recognition of other bacterial effectors (*P. syringae* HopZ5, AvrRpm1, AvrB and *X. euvesicatoria* AvrBsT) that are sequence-unrelated to AvrRpt2 but known to modify RIN4 (Ahn et al., 2022; Choi et al., 2021; Chung et al., 2011; Liu et al., 2011).

Here, we show that *NbPtr1* is necessary and sufficient for RipE1 recognition using a reverse-genetic approach based on multiplexed silencing of *N. benthamiana NLR* genes. We also demonstrate that RipE1 recognition depends on its association with the host cell plasma membrane. Lastly, our survey of RipE1 allelic diversity across the RSSC supports the broad conservation of NbPtr1-dependent recognition, providing an additional evidence for the indirect mode of activation of this NLR.

To identify NLR(s) involved in RipE1 recognition, we screened a recently developed NLR-VIGS library (Ahn et al., 2022) for loss of RipE1-induced cell death. RipE1-induced cell death was completely abolished in plants silenced with the TRV:Com49 construct which targets several ungrouped *CNLs*, and more specifically in plants silenced with the TRV:Com49-3 that targets only *NbPtr1* (Ahn et al., 2022) (Figure S1). To confirm *NbPtr1* involvement in RipE1-induced cell death, we used two alternative VIGS fragments for specific silencing of *NbPtr1* (Figure 1a). While RipE1 expression led to a robust macroscopic cell death in TRV:EV plants, it failed to induce cell death in both TRV:*NbPtr1*.*a* and TRV:*NbPtr1*.*b* plants. The cell death intensity was quantified using the chlorophyll quantum yield value in leaf tissue expressing RipE1 or the unrecognized RipE1-C172A mutant. In accordance, RipE1-expressing leaf tissue displayed a higher QY in TRV:*NbPtr1*.*a* and TRV:*NbPtr1*.*b* plants compared to the TRV:EV control plants (Figure 1b). *NbPtr1* silencing efficiency was confirmed by qRT-PCR in repeated experiments, as well as the stability of RipE1 proteins in the examined tissues (Figures S2a and S2b).

**Figure 1.**
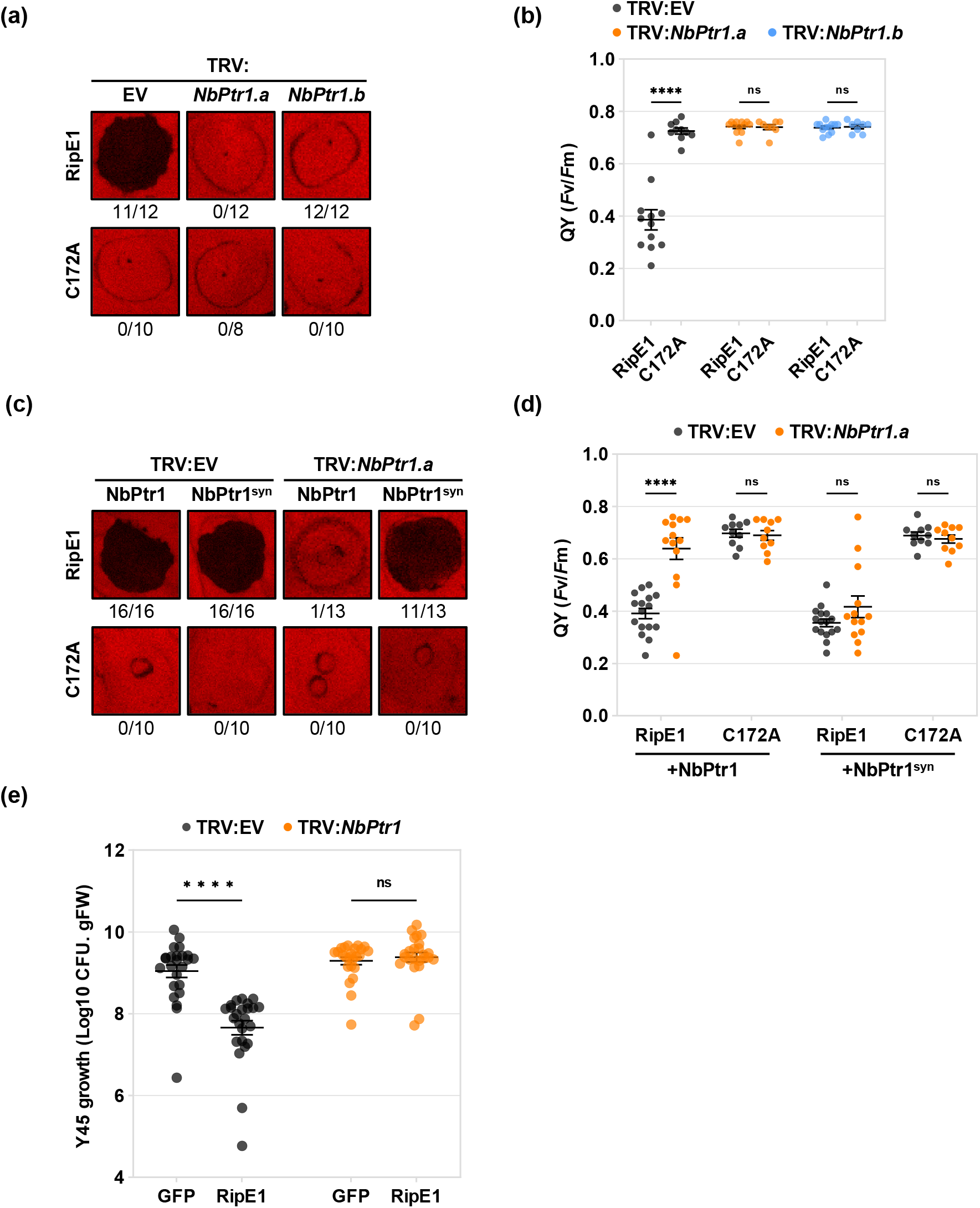
*NbPtr1* is genetically required for RipE1 recognition in *N. benthamiana*. (a) RipE1-induced cell death was suppressed in *NbPtr1*-silenced plants. Agrobacteria carrying RipE1 or RipE1-C172A constructs were infiltrated (OD_600_ = 0.5) in TRV:EV, TRV:*NbPtr1*.*a* and TRV:*NbPtr1*.*b* plants. Photographs were taken 4 days post-infiltration (dpi) under LED light (false color scale). Numbers indicate patches with cell death out of total infiltrated patches. (b) Quantum yield of the photosystem II (QY, *F*v/*F*m) was measured in the infiltrated patches shown in (a). Individual values from independent experiments are indicated as dots. Data were analyzed with two-way ANOVA followed by Sidak’s multiple comparison test. Asterisks indicate significant difference between RipE1 constructs in TRV:EV, TRV:*NbPtr1*.*a* and TRV:*NbPtr1*.*b* plants (****, *P*<0.0001; ns, not significant). Bars represent mean ± SEM (n=8-12). (c) Silencing-proof *NbPtr1* (NbPtr1^syn^) expression restored RipE1-induced cell death in *NbPtr1*-silenced plants. NbPtr1 or NbPtr1^syn^ (OD_600_ = 0.05) were co-expressed with RipE1 or RipE1-C172A (OD_600_ = 0.4) in TRV:EV and TRV:*NbPtr1*.*a* plants. Photographs were taken at 4 dpi under LED light. Numbers indicate patches with cell death out of total infiltrated patches. (d) Quantum yield of the photosystem II (QY, *F*v/*F*m) was measured in the infiltrated patches shown in (c). Data were analyzed with two-way ANOVA followed by Sidak’s multiple comparison test. Asterisks indicate significant difference between RipE1 constructs in TRV:EV and TRV:*NbPtr1*.*a* plants (****, *P*<0.0001; ns, not significant). (e) RipE1-induced resistance to *R. pseudosolanacearum* was suppressed in *NbPtr1*-silenced plants. *R. pseudosolanacearum* Y45 was infiltrated at 10^5^ CFU. ml^-1^ in TRV:EV and TRV:*NbPtr1*.*a* plants one day after agroinfiltration of GFP or RipE1-GFP (OD_600_ = 0.2). bacteria were enumerated at 2 dpi. Individual values from independent experiments are indicated as dots. Data were analyzed with two-way ANOVA followed by Sidak’s multiple comparison test. Asterisks indicate significant difference with GFP in TRV:EV and TRV:*NbPtr1*.*a* plants (****, *P*<0.0001; ns, not significant). Bars represent mean ± SEM (n=24).

In order to additionally confirm the NbPtr1 function in RipE1 recognition, we next performed a genetic complementation assay using a synthetic *NbPtr1* construct containing alternative synonymous codons at the beginning of the first exon to prevent silencing triggered by the *NbPtr1*.*a* fragment (Ahn et al., 2022; Wu et al., 2017) (Figure 1c). RipE1 or RipE1-C172A mutant were co-expressed with the full-length NbPtr1 or the silencing-proof NbPtr1^syn^ in both TRV:EV and TRV:*NbPtr1*.*a* plants. NbPtr1^syn^ co-expression with RipE1 could restore a robust cell death in TRV:*NbPtr1*.*a* plants and this correlated well with a low QY (Figure 1d). We confirmed the stability of NbPtr1^syn^ and the lack of accumulation of NbPtr1 protein in the TRV:*NbPtr1*.*a* plants by immunoblotting (Figure S2c). Similar assays were further conducted in Nb*-ptr1* knockout plants, where the *NbPtr1* coding sequence is disrupted by a CRISPR-induced 61 bp deletion (Ahn et al., 2022) (Figure S3). RipE1-induced cell death was completely abolished in Nb*-ptr1* plants and could be restored by expression of the native *NbPtr1* gene, demonstrating that *NbPtr1* is genetically necessary and sufficient to induce cell death in presence of RipE1 in *N. benthamiana*.

We further investigated the role of *NbPtr1* in RipE1-triggered immunity to *R. pseudosolanacearum*. We used the virulent *R. pseudosolanacearum* Y45 strain that lacks several effectors recognized by the *N. benthamiana* immune system, including RipAA, RipP1 and RipE1 (Li et al., 2011; Pouymero et al., 2009; Sang et al., 2020; Yu and Macho, 2021). *N. benthamiana* plants silenced with TRV:EV or TRV:*NbPtr1*.*a* were infected with the Y45 strain one day after infiltration with Agrobacterium strains for the expression of RipE1-GFP or GFP (Figure 1e). Consistent with the previous report (Sang et al., 2020), Y45 cells grew ∼10 times less in TRV:EV plants expressing RipE1 compared to TRV:EV plants expressing GFP. This growth difference was strikingly abolished in TRV:*NbPtr1*.*a* plants, where Y45 grew to a similar level in both GFP-and RipE1-expressing tissues. As we confirmed the stability of RipE1-GFP protein in this assay (Figure S2d), these results demonstrate that *NbPtr1* mediates the recognition of RipE1 and contributes to RipE1-triggered immunity to *R. pseudosolanacearum*. It is challenging to dissect the genetic basis of resistance to bacterial wilt disease, as a large number of effectors are concomitantly delivered, and elicit or prevent the plant immune responses. It is noteworthy that two other *R. pseudosolanacearum* effectors, RipAY and RipAC, are suppressors of the RipE1-induced immune responses in *N. benthamiana* (Sang et al., 2020; Yu et al., 2020). Nonetheless, a reverse genetic approach and heterologous expression of effectors allowed us to rapidly identify avirulence effector/immune receptor pairs, expanding the pool of interactions to be considered for crop improvement.

As NbPtr1 is required for the recognition of effectors from different families, likely by monitoring the state of the plasma membrane-associated protein RIN4 (Ahn et al., 2022; Mazo-Molina et al., 2020), we next investigated RipE1 subcellular localization with laser scanning confocal microscopy. Nakano and colleagues (2019) reported a nucleocytoplasmic localization for the N-terminally tagged RipE1 (from strain RS1000), while we and others (Jeon et al., 2020; Tsakiri et al., 2022) observed a plasma membrane localization for RipE1 C-terminally tagged with YFP. This difference suggests that RipE1 association with the plasma membrane is mediated through the N-terminal region of the protein. Accordingly, we expressed C-terminally YFP-tagged RipE1, RipE1-C172A and RipE1-ΔN86, a truncated variant lacking the first N-terminal 86 amino acids in Nb*-ptr1* epidermal cells (Figure 2a). RipE1-YFP was clearly associated with the plasma membrane, as the YFP fluorescence overlapped with that of the plasma membrane marker AtFLS2-mCherry. Interestingly, while RipE1-C172A-YFP remained in association with the plasma membrane, the RipE1-ΔN86-YFP presented a nucleocytoplasmic localization. Moreover, similar to RipE1-C172A, RipE1-ΔN86 could not induce cell death in *N. benthamiana*, despite showing a comparable protein stability (Figure 2b, 2c and 2d). These findings indicate that RipE1 association with the plasma membrane is required for RipE1-induced cell death in addition to the putative catalytic activity. The N-terminal region of RipE1 was not predicted to participate in the cysteine-protease fold in a predicted structural model (Tsakiri et al., 2022). This region does not contain a myristylation motif that was reported for several plasma membrane-associated T3S effectors including AvrRpm1 (Nimchuk et al., 2000). However, RipE1 could be S-acylated in this region for its recruitment to the plasma membrane after delivery into the host cell (Hurst and Hemsley, 2015).

**Figure 2.**
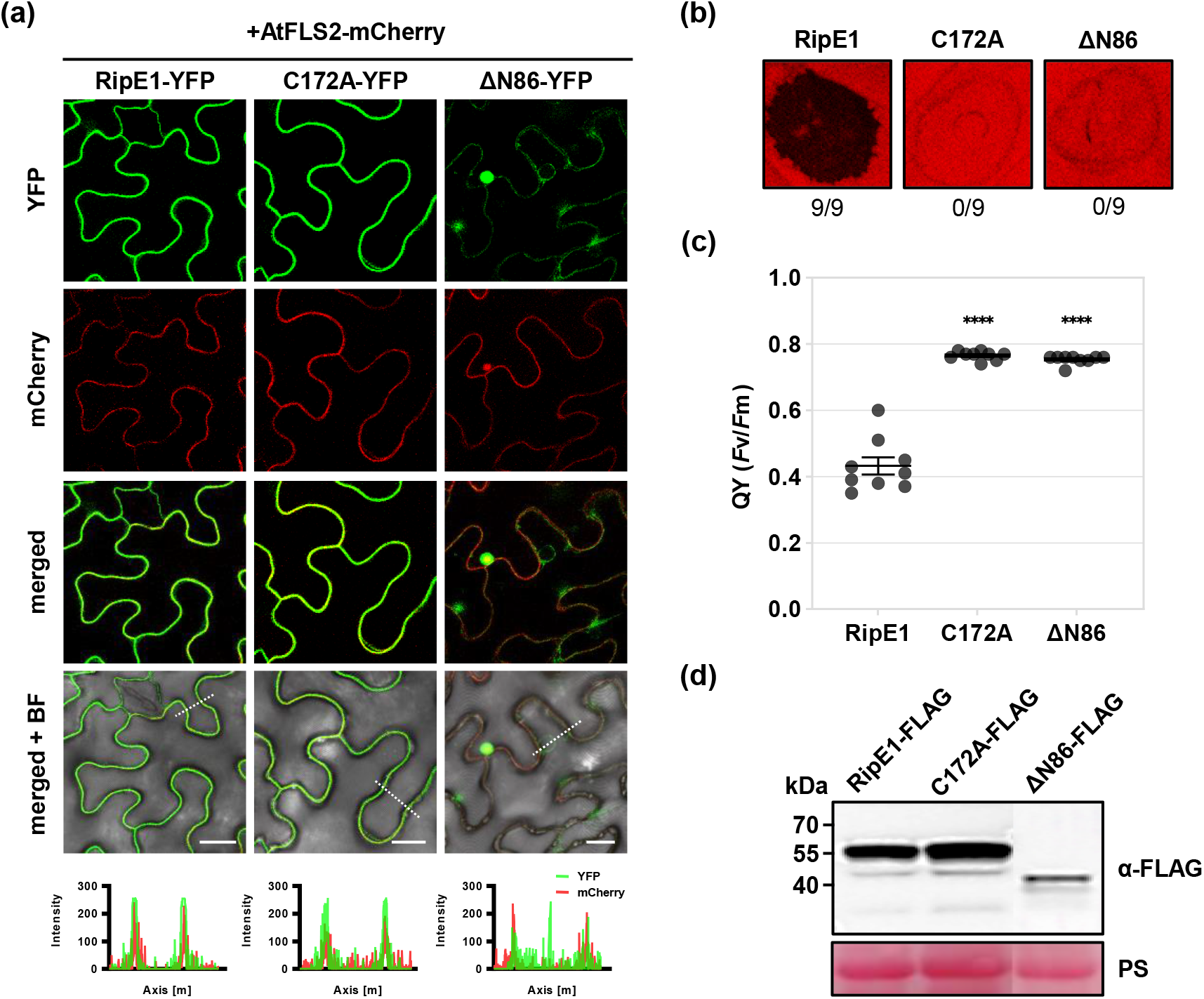
RipE1 association with the plasma membrane is essential for recognition by *NbPtr1*. (a) RipE1 N-terminus is required for association with the plasma membrane. RipE1, RipE1-C172A and RipE1-ΔN86 C-terminally fused with YFP (OD_600_ = 0.4) were co-expressed with the plasma membrane marker AtFLS2-mCherry (OD_600_ = 0.1) in Nb*-ptr1* epidermal cells. Confocal micrographs were acquired at 2 dpi (BF, bright field). The fluorescence intensity of YFP and mCherry channels across sections indicated by dotted lines are shown in the bottom panel. Scale bar indicates 11 µm. (b) RipE1 N-terminus is required for recognition by NbPtr1. Agrobacteria carrying RipE1, RipE1-C172A or RipE1-ΔN86 were infiltrated (OD_600_ = 0.4) in WT plants. Photographs were taken at 4 dpi under LED light. Numbers indicate patches with cell death out of total infiltrated patches. (c) Quantum yield of the photosystem II (QY, *F*v/*F*m) was measured in the infiltrated patches shown in (b). Individual values from independent experiments are indicated as dots. Data were analyzed with one-way ANOVA followed by Dunnett’s multiple comparison test. Asterisks indicate significant difference with RipE1 (****, *P*<0.0001; ns, not significant). Bars represent mean ± SEM (n=9). (d) RipE1, RipE1-C172A and RipE1-ΔN86 accumulate in WT plants. Immunodetection was performed with anti-FLAG antibodies on total protein extracted at 36 h post-infiltration. Ponceau red staining (PS) shows equal loading of the samples.

Lastly, we investigated the natural variation of RipE1 alleles and the specificity of the recognition by *NbPtr1*. RipE1 is present in all the strains representative of the diversity of *Ralstonia* spp. (Peeters et al., 2013; Sabbagh et al., 2019). We retrieved 48 unique RipE1 sequences that could be grouped into three clades corresponding to *R. pseudosolanacearum, R. solanacearum* and *R. syzygii* strains (Figure S4a). Of note, several *R. syzygii* strains isolated from tomato carry two alleles of RipE1, one being more distant from all the other natural variants. We selected and synthesized RipE1 sequences present in strains CMR15 (*R. pseudosolanacearum*), Po82 (*R. solanacearum*) and PSI07 (*R. syzygii*) to evaluate the impact of natural variation on the recognition by *NbPtr1*. These alleles present 87%, 84% and 42% amino acid identity, respectively, with RipE1_GMI1000 used in all our previous experiments but they all harbor the conserved catalytic triad residues and the conserved domain A, previously reported as required for the recognition by plant immune system (Nimchuk et al., 2007; Sang et al., 2020) (Figure S4b). As observed with RipE1_GMI1000, RipE1_CMR15 and RipE1_Po82 expression induced a robust cell death correlated with significantly low QY in WT plants (Figure 3a and 3b). This response was abolished in Nb*-ptr1* plants and restored by co-expression with NbPtr1. Interestingly, RipE1_PSI07 variant induced a significantly weak cell death in WT plants but co-expression with *NbPtr1* restored a robust cell death in Nb*-ptr1* plants. All transiently expressed RipE1 variants used in our study accumulated to a similar level indicating that the reduced cell death observed with RipE1_PSI07 protein was not due to its unstable nature (Figure 3c). Considering the importance of RipE1 subcellular localization for its recognition by NbPtr1, we examined the localization of the selected RipE1 variants C-terminally fused with YFP (Figure S5). The YFP signal overlapped with the AtFLS2-mCherry marker indicated that RipE1_PSI07, similar to RipE1_CMR15 and RipE1_Po82, could be associated with the plasma membrane. These results suggest that RipE1_PSI07 might lack additional features contributing to NbPtr1 activation that could be compensated by the overexpression of *NbPtr1* in our experimental conditions.

**Figure 3.**
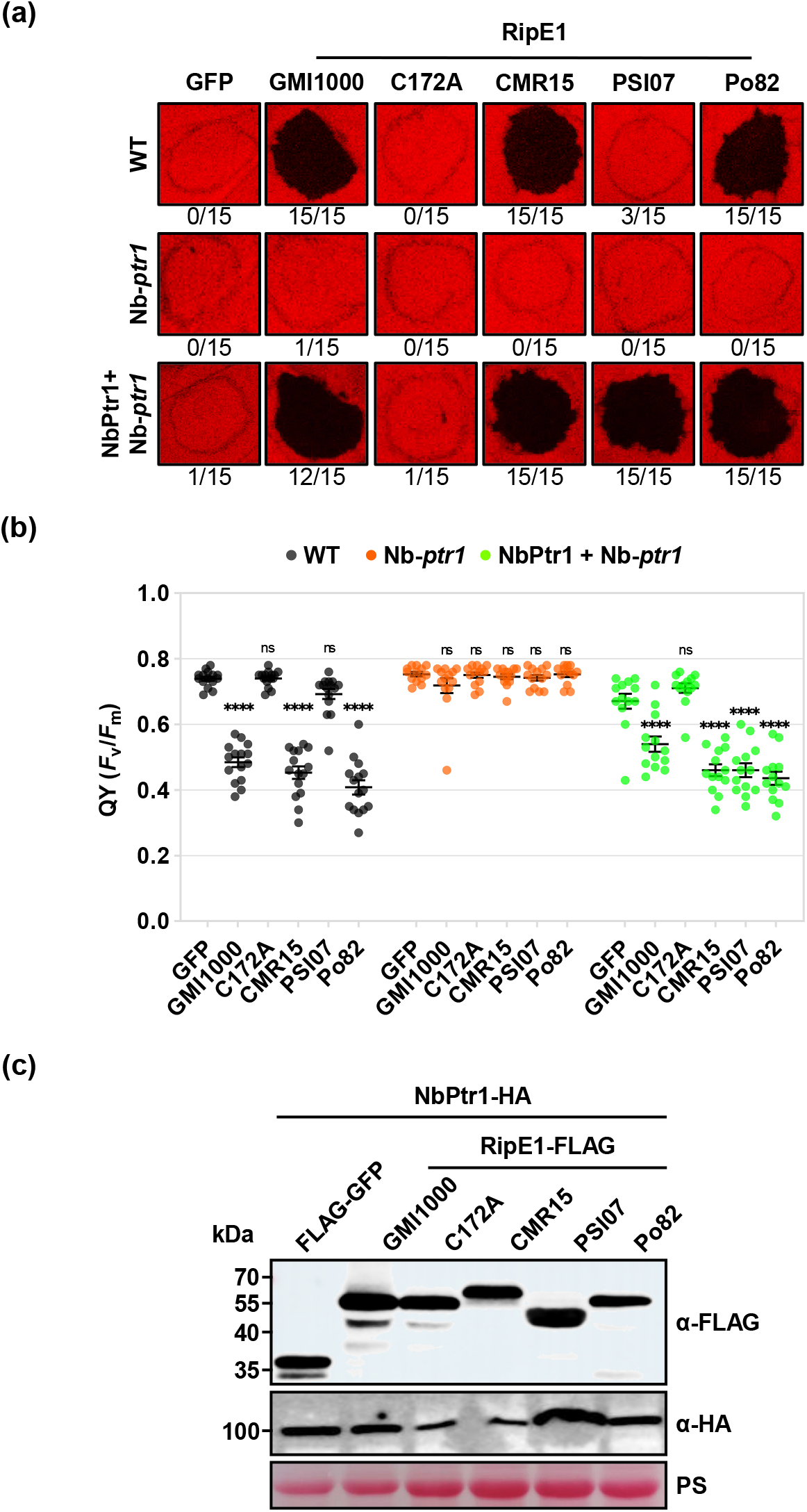
NbPtr1 recognizes RipE1 natural variants from different *Ralstonia* species. (a) RipE1 natural variants induced *NbPtr1*-dependent cell death. Agrobacteria carrying GFP, RipE1 (GMI1000), RipE1-C172A, or RipE1 variants from *R. pseudosolanacearum* strain CMR15 (CMR15), *R. solanacearum* strain Po82 (Po82) and *R. syzygii* strain PSI07 (PSI07_1845) constructs were infiltrated (OD_600_ = 0.4) in WT or Nb*-ptr1* plants without or with NbPtr1 (OD_600_ = 0.05). Photographs were taken at 4 dpi under LED light. Numbers indicate patches with cell death out of total infiltrated patches. (b) Quantum yield of the photosystem II (QY, *F*v/*F*m) was measured in the infiltrated patches shown in (a). Individual values from independent experiments are indicated as dots. Data were analyzed with two-way ANOVA followed by Dunnett’s multiple comparison test. Asterisks indicate significant difference with GFP (****, *P*<0.0001; ns, not significant). Bars represent mean ± SEM (n=15). (c) RipE1 natural variants and NbPtr1 accumulate in Nb*-ptr1* plants. Immunodetection was performed with anti-FLAG and anti-HA antibodies on total protein extracted at 36 h post-infiltration. Ponceau red staining (PS) shows equal loading of the samples.

In summary, our data demonstrate that NbPtr1 can recognize the RipE1 natural variants originating from multiple *Ralstonia* species. This suggests that *NbPtr1* may be a valuable resource for developing a durable bacterial wilt-resistant crops. Further, the importance of the plasma membrane for RipE1 function will help better understand the detailed activation mechanism of NbPtr1 in the future.

## Author contribution

Conceptualization: CS, KHS; Investigation: BK, IK, WY, HK, YJA; Resources: CS, KHS, APM; Supervision: CS; Funding acquisition: BK, CS; Writing: CS with inputs from all the authors.

## Acknowledgments

We would like to thank Ning Zhang and Gregory Martin for kindly providing the Nb*-ptr1* line. This work was supported by the National Research Foundation of Korea (NRF) funded by the Korean Ministry of Education (Global Ph.D. Fellowship Program Project No 500-20190213) and by the Ministry of Sciences and ICT (Projects No 2018R1A5A1023599 and No 2020R1A2C1101419). The authors declare no conflict of interest.

## Data availability statement

The data that support the findings of this study are available from the corresponding author upon reasonable request.

## Supporting Information Legends

Figure S1. RipE1-induced cell death is abolished in TRV:Com49-3 silenced plants. (a) Agrobacteria carrying GFP, RipE1, or RipE1-C172A (OD_600_ = 0.5) constructs were infiltrated in TRV:EV, TRV:Com-49, and TRV:Com-49-1 to -6 plants. Photographs were taken 3 days post-infiltration (dpi) under LED light (false color scale). Numbers indicate patches with cell death out of total infiltrated patches. (b) Quantum yield of the photosystem II (QY, *F*v/*F*m) was measured in the infiltrated patches shown in (a). Data were analyzed with two-way ANOVA followed by Dunnett’s multiple comparison test. Asterisks indicate significant difference with GFP (****, *P*<0.0001). Bars represent mean ± SEM (n=3-6).

Figure S2. Analysis of *NbPtr1* silencing and RipE1 accumulation in TRV:*NbPtr1* plants. (a) *NbPtr1* expression was measured by qRT-PCR in TRV:EV, TRV:*NbPtr1*.*a* and TRV:*NbPtr1*.*b* plants and normalized to *NbEF1α* expression. Individual values from independent experiments are indicated as dots. Data were analyzed with one-way ANOVA followed by Dunnett’s multiple comparison test. Asterisks indicate significant difference with TRV:EV plants (****, *P*<0.0001). Bars represent mean ± SEM (n=15). (b) RipE1 and RipE1-C172A C-terminally fused to 3xFLAG tag accumulate in *NbPtr1*-silenced plants. Immunodetection was performed with anti-FLAG antibodies on total protein extracted at 36 h post-infiltration (hpi). Ponceau red staining (PS) attests equal loading of the samples. (c) NbPtr1^syn^ accumulates in TRV:*NbPtr1* plants. Immunodetection was performed with anti-HA antibodies on total proteins extracted at 36 hpi. (d) RipE1-GFP accumulates in TRV:*NbPtr1* plants. Immunodetection was performed with anti-GFP antibodies on total protein extracts. Coomassie blue staining (CBB) attests equal loading of the samples.

Figure S3. RipE1 recognition is abolished in Nb*-ptr1* knock-out plants. (a) Native NbPtr1 is sufficient to restore RipE1-induced cell death in Nb*-ptr1* plants. Agrobacteria carrying RipE1, RipE1-C172A (OD_600_ = 0.4) or NbPtr1 (OD_600_ = 0.05) constructs were infiltrated in WT or *Nb-ptr1* plants. Photographs were taken 4 days post-infiltration (dpi) under LED light (false color scale). Numbers indicate patches with cell death out of total infiltrated patches. (b) Quantum yield of the photosystem II (QY, *F*v/*F*m) was measured in the infiltrated patches shown in (a). Individual values from independent experiments are indicated as dots. Data were analyzed with two-way ANOVA followed by Sidak’s multiple comparison test. Asterisks indicate significant difference with WT plants (****, *P*<0.0001; ns, not significant). Bars represent mean ± SEM (n=8-16). (c) RipE1 and NbPtr1 accumulate in WT and Nb*-ptr1* plants. Immunodetection was performed with anti-FLAG or anti-HA antibodies on total protein extracted at 36 hpi. Ponceau red staining (PS) attests equal loading of the samples.

Figure S4. RipE1 natural variation across *Ralstonia* species. (a) Phylogenetic tree of 48 unique RipE1 proteins from sequenced *Ralstonia* spp strains. RipE1 protein sequences were obtained from Ralsto T3E database (htpps://iant.toulouse.inra.fr/bacteria/annotation/site/prj/T3Ev3/). The tree was built using Jukes-Cantor genetic distance model with the neighbor-joining method and the *R. pseudosolanacearum* RipBN effector as the outgroup. RipE1 sequences highlighted in bold were cloned for transient expression in *N. benthamiana*. (b) Protein sequence alignment of the RipE1 variants selected in this study. Black and grey highlight amino acid identity. The predicted domain A and the Cys, His, Asp catalytic triad are indicated in blue.

Figure S5. RipE1 natural variants associate with the plasma membrane. (a) RipE1 from strain CMR15, Po82 and PSI07 C-terminally fused with YFP (OD_600_ = 0.4) were co-expressed with the plasma membrane marker AtFLS2-mCherry (OD_600_ = 0.1) in Nb*-ptr1* epidermal cells. Confocal micrographs were acquired at 2 dpi (BF, bright field). The fluorescence intensity of YFP and mCherry channels across sections indicated by dotted lines are shown in the bottom panel. Scale bar indicates 11 µm. (b) Accumulation of RipE1-YFP fusion proteins in nbptr1 plants. Immunodetection was performed with anti-GFP or anti-mCherry antibodies on total protein extracted at 36 hpi. Ponceau red staining (PS) attests equal loading of the samples.

