## Supplemental Figures for "The *Ralstonia pseudosolanacearum* effector RipE1 is recognized at the plasma membrane by *NbPtr1, Nicotiana benthamiana* homolog of *Pseudomonas tomato race 1*"

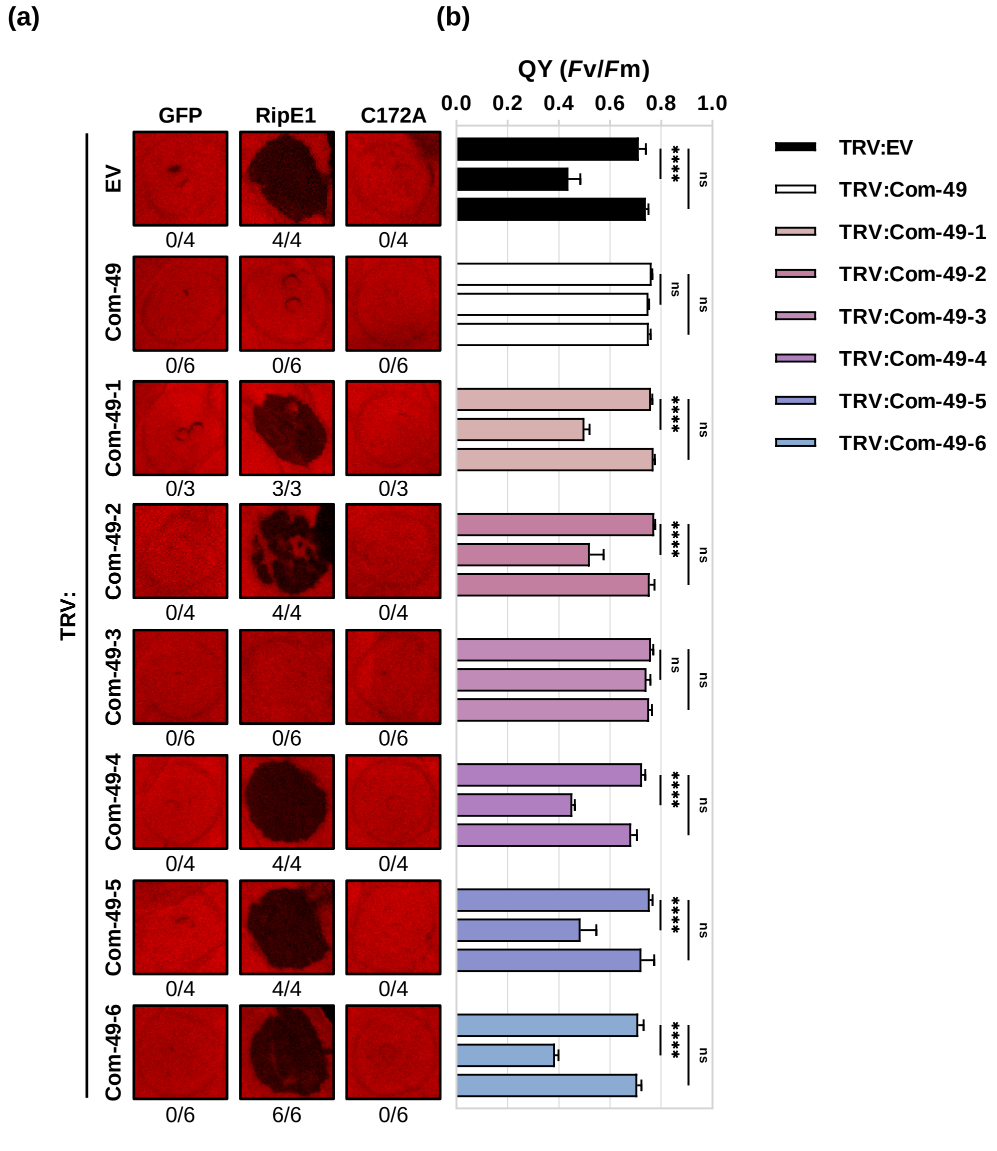


Figure S1. RipE1-induced cell death is abolished in TRV:Com49-3 silenced plants. (a) Agrobacteria carrying GFP, RipE1, or RipE1-C172A ($OD_{600}$ = 0.5) constructs were infiltrated in TRV:EV, TRV:Com-49, and TRV:Com-49-1 to -6 plants. Photographs were taken 3 days post-infiltration (dpi) under LED light (false color scale). Numbers indicate patches with cell death out of total infiltrated patches. (b) Quantum yield of the photosystem II (QY, *Fv*/*Fm*) was measured in the infiltrated patches shown in (a). Data were analyzed with two-way ANOVA followed by Dunnett's multiple comparison test. Asterisks indicate significant difference with GFP (****, *P*<0.0001; ns, not significant). Bars represent mean ± SEM (n=3-6).

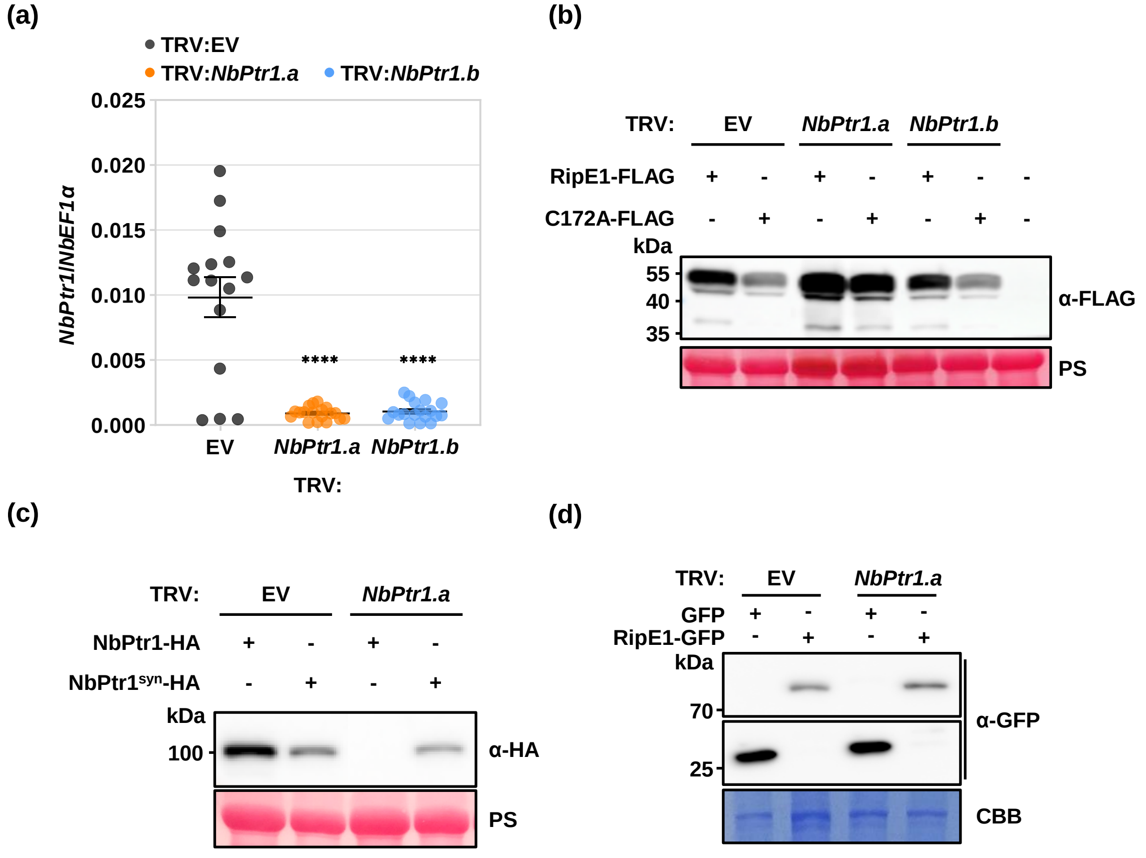


Figure S2. Analysis of *NbPtr1* silencing and RipE1 accumulation in TRV:*NbPtr1* plants. (a) *NbPtr1* expression was measured by qRT-PCR in TRV:EV, TRV:*NbPtr1.a* and TRV:*NbPtr1.b* plants and normalized to *NbEF1α* expression. Individual values from independent experiments are indicated as dots. Data were analyzed with one-way ANOVA followed by Dunnett's multiple comparison test. Asterisks indicate significant difference with TRV:EV plants (****, *P*<0.0001). Bars represent mean ± SEM (n=15). (b) RipE1 and RipE1-C172A C-terminally fused to 3xFLAG tag accumulate in *NbPtr1*-silenced plants. Immunodetection was performed with anti-FLAG antibodies on total protein extracted at 36 h post-infiltration (hpi). Ponceau red staining (PS) attests equal loading of the samples. (c) NbPtr1$^{syn}$ accumulates in TRV:*NbPtr1.a* plants. Immunodetection was performed with anti-HA antibodies on total proteins extracted at 36 hpi. (d) RipE1-GFP accumulates in TRV:*NbPtr1.a* plants. Immunodetection was performed with anti-GFP antibodies on total protein extracts. Coomassie blue staining (CBB) attests equal loading of the samples.

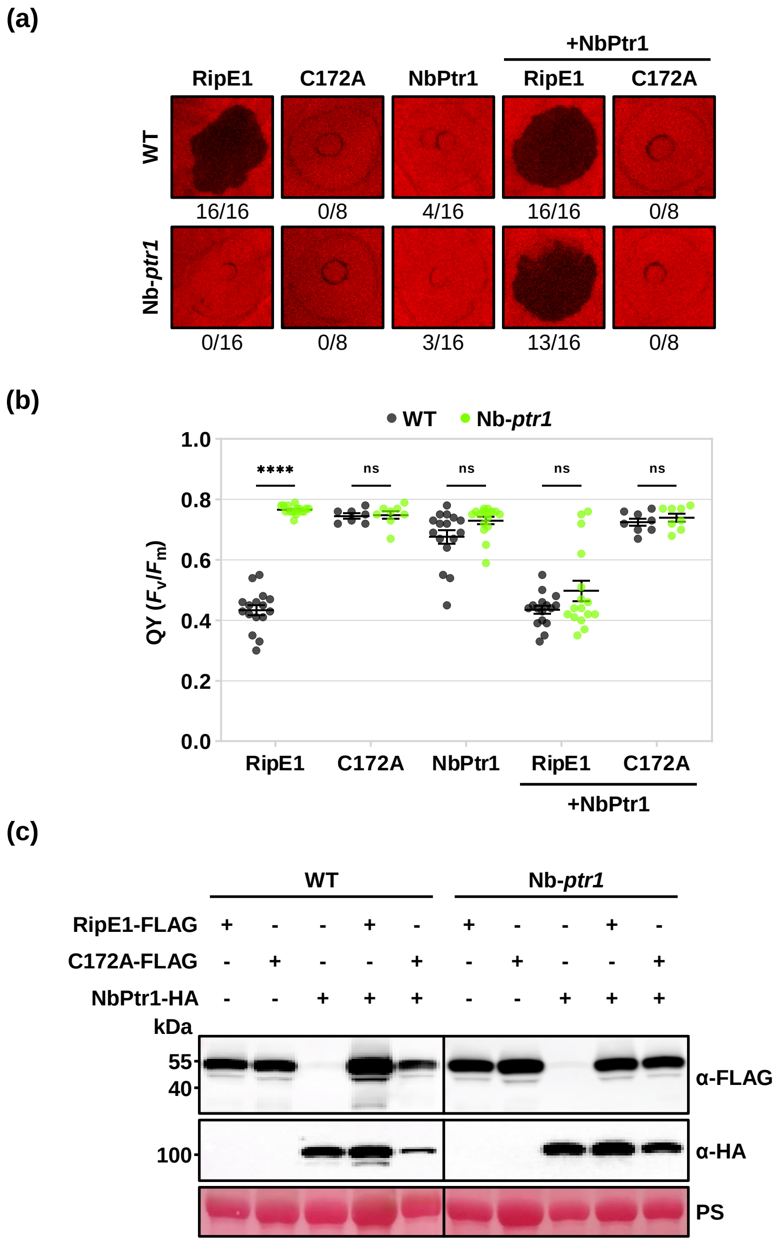


Figure S3. RipE1 recognition is abolished in Nb-*ptr1* knock-out plants. (a) Native NbPtr1 is sufficient to restore RipE1-induced cell death in Nb-*ptr1* plants. Agrobacteria carrying RipE1, RipE1-C172A ($OD_{600}$ = 0.4) or NbPtr1 ($OD_{600}$ = 0.05) constructs were infiltrated in WT or Nb-*ptr1* plants. Photographs were taken at 4 dpi under LED light. Numbers indicate patches with cell death out of total infiltrated patches. (b) Quantum yield of the photosystem II (QY, *Fv*/*F*m) was measured in the infiltrated patches shown in (a). Individual values from independent experiments are indicated as dots. Data were analyzed with two-way ANOVA followed by Sidak's multiple comparison test. Asterisks indicate significant difference with WT plants (****, $P<0.0001$; ns, not significant). Bars represent mean ± SEM (n=8-16). (c) RipE1 and NbPtr1 accumulate in WT and Nb-*ptr1* plants. Immunodetection was performed with anti-FLAG or anti-HA antibodies on total protein extracted at 36 h post-infiltration (hpi). Ponceau red staining (PS) attests equal loading of the samples.

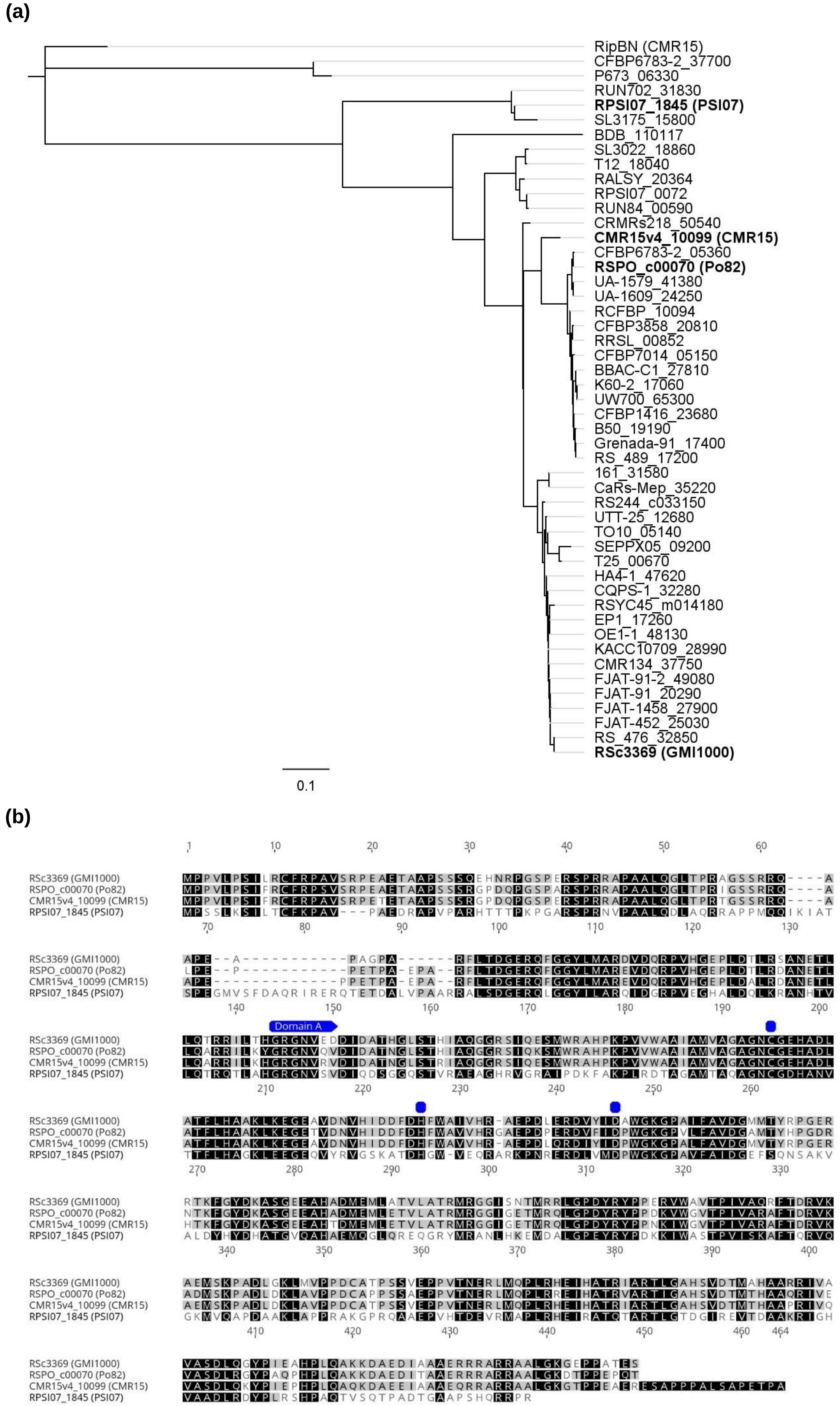


Figure S4. RipE1 natural variation across *Ralstonia* species. (a) Phylogenetic tree of 48 unique RipE1 proteins from sequenced *Ralstonia* spp. strains. RipE1 protein sequences were obtained from Ralsto T3E database (htpps://iant.toulouse.inra.fr/bacteria/annotation/site/prj/T3Ev3/). The tree was built using Jukes-Cantor genetic distance model with the neighbor-joining method and the *R. pseudosolanacearum* RipBN effector as the outgroup. RipE1 sequences highlighted in bold were cloned for transient expression in *N. benthamiana*. (b) Protein sequence alignment of the RipE1 variants selected in this study. Black and grey highlight amino acid identity. The predicted domain A and the Cys, His, Asp catalytic triad are indicated in blue.

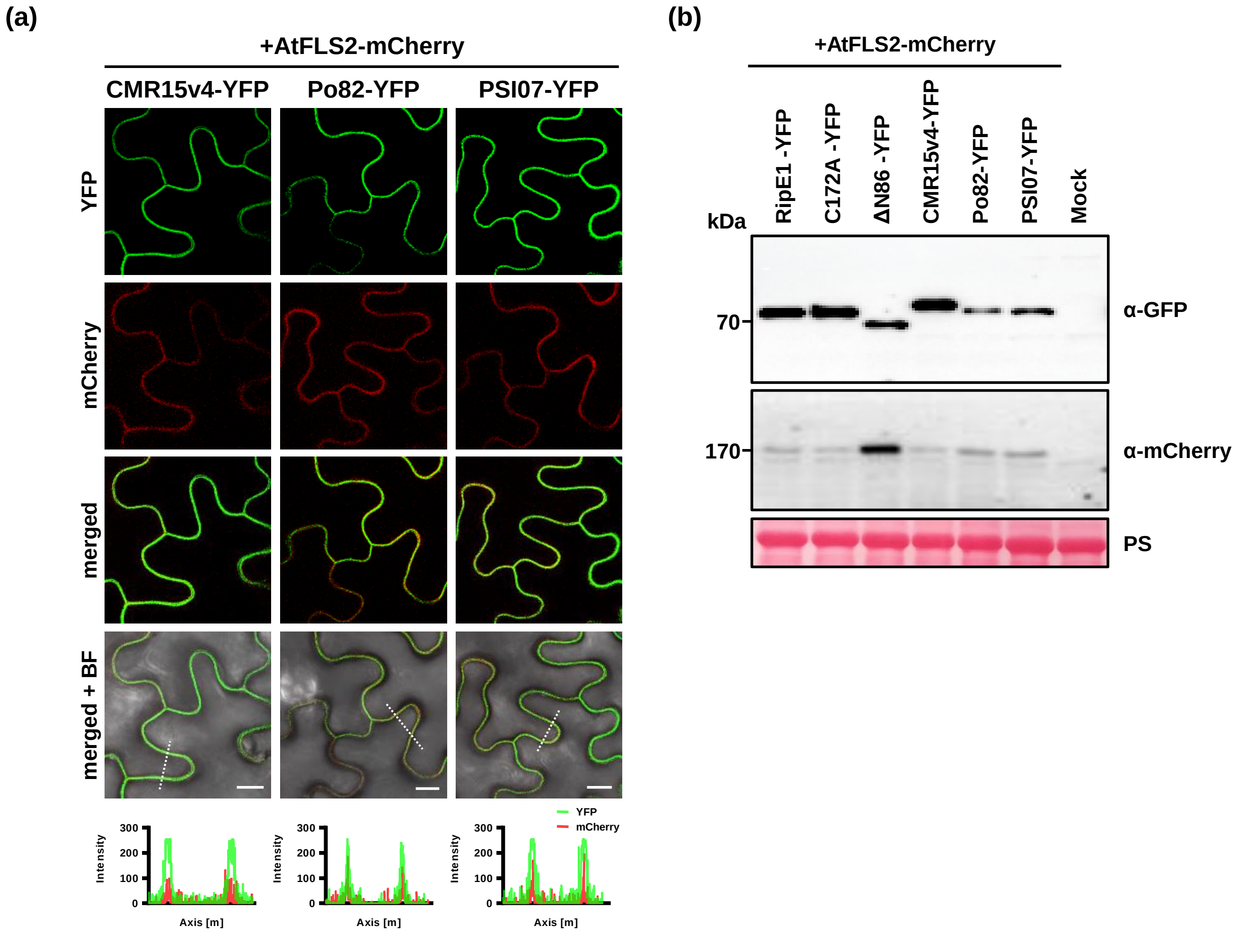


Figure S5. RipE1 natural variants associate with the plasma membrane. (a) RipE1 from strain CMR15, Po82 and PSI07 C-terminally fused with YFP ($OD_{600}$ = 0.4) were co-expressed with the plasma membrane marker AtFLS2-mCherry ($OD_{600}$ = 0.1) in Nb-*ptr1* epidermal cells. Confocal micrographs were acquired at 2 dpi (BF, bright field). The fluorescence intensity of YFP and mCherry channels across sections indicated by dotted lines are shown in the bottom panel. Scale bar indicates 11 μm. (b) Accumulation of RipE1-YFP fusion proteins in Nb-*ptr1* plants. Immunodetection was performed with anti-GFP or anti-mCherry antibodies on total protein extracted at 36 hpi. Ponceau red staining (PS) attests equal loading of the samples.
